# Defining the Polycystin Pharmacophore Through HTS & Computational Biophysics

**DOI:** 10.1101/2025.01.13.632808

**Authors:** Eduardo Guadarrama, Carlos G. Vanoye, Paul G. DeCaen

## Abstract

**Background and Purpose:** Polycystins (PKD2, PKD2L1) are voltage-gated and Ca^2+^-modulated members of the transient receptor potential (TRP) family of ion channels. Loss of PKD2L1 expression results in seizure-susceptibility and autism-like features in mice, whereas variants in PKD2 cause autosomal dominant polycystic kidney disease. Despite decades of evidence clearly linking their dysfunction to human disease and demonstrating their physiological importance in the brain and kidneys, the polycystin pharmacophore remains undefined. Contributing to this knowledge gap is their resistance to drug screening campaigns, which are hindered by these channels’ unique subcellular trafficking to organelles such as the primary cilium. PKD2L1 is the only member of the polycystin family to form constitutively active ion channels on the plasma membrane when overexpressed.

**Experimental Approach:** HEK293 cells stably expressing PKD2L1 F514A were pharmacologically screened via high-throughput electrophysiology to identify potent polycystin channel modulators. In-silico docking analysis and mutagenesis were used to define the receptor sites of screen hits. Inhibition by membrane-impermeable QX-314 was used to evaluate PKD2L1’s binding site accessibility.

**Key Results:** Screen results identify potent PKD2L1 antagonists with divergent chemical core structures and highlight striking similarities between the molecular pharmacology of PKD2L1 and voltage-gated sodium channels. Docking analysis, channel mutagenesis, and physiological recordings identify an open-state accessible lateral fenestration receptor within the pore, and a mechanism of inhibition that stabilizes the PKD2L1 inactivated state.

**Conclusion and Implication:** Outcomes establish the suitability of our approach to expand our chemical knowledge of polycystins and delineates novel receptor moieties for the development of channel-specific antagonists in TRP channel research.

## Regarding polycystin nomenclature

The revised and current IUPHAR/BPS nomenclature creates ambiguity regarding the genetic identity of the polycystin family members of transient receptor potential ion channels (TRPP), especially when cross-referencing manuscripts that describe subunits using the former system.^1^ Traditionally, the products of polycystin genes (e.g., PKD2) are referred to as polycystin proteins (e.g., polycystin-2). For simplicity and to prevent confusion, we will refer to the polycystin gene name rather than differentiating gene and protein with separate names— a nomenclature we have recently outlined.^2^

## INTRODUCTION

Transient receptor potential (TRP) cation channels function as polymodal cellular sensors that open in response to a broad range of chemical, thermal and physical stimuli— resulting in electrical depolarization of nerves and other cell membranes^3–5^. Humans express a total of 28 different TRP channel proteins, which can be divided into seven subfamilies based on amino acid sequence homology: ankyrin, canonical, melastatin, mucolipin, polycystin and vanilloid^6,7^. Members of the polycystin class (PKD2, PKD2L1, PKD2L2) have six transmembrane segments which can assemble as homotetrameric channels that are voltage-gated and Ca^2+^-modulated^8–13^. Polycystin subunits also form heteromeric complexes with closely-related, eleven-transmembrane-segment proteins (PKD1, PKD1L1, PKD1L2), which contain both channel and adhesion g-protein-coupled-receptor components^2,14,15^. Heteromeric polycystin channels are proposed to be cleavage or ligand activated, but their native functional properties are still being elucidated^16,17^. Several members of the polycystin family show temporal expression during embryonic development (e.g. embryonic node) and localize to divergent organ tissues in adulthood, such as kidney principle cells and hippocampal neurons^18,19^. Human clinical genetics and mouse transgenic studies indicate that polycystins are critically involved in disparate forms of cell physiology and function as “organizers” of cell differentiation during gastrulation^20,21^. Human germline variants in renal polycystins (PKD1 and PKD2) are responsible for autosomal dominant polycystic kidney disease (ADPKD), a monogenetic disorder affecting ≈13 million individuals, characterized by the proliferation of fluid-filled cysts that precipitate renal failure^22–24^. In contrast, PKD2L1 variants are not reportedly associated with human disease, but loss of their expression causes hippocampal neuron hyperexcitability, which precipitates seizure susceptibility and autism spectrum disorder-like neurophenotypes in mice^19,25,26^.

Despite their clear association with human disease and their roles in human development and cell physiology, the pharmacology of polycystins is largely undescribed. This is principally due to their unique subcellular localization, which makes them resistant to drug screening. Renal polycystins (PKD2) traffic to, and form functional channels in, the membranes of primary cilia organelles in kidney principal cells and when overexpressed in cell lines^9–11^. Importantly, they are electrically and chemically insulated from the cytosolic compartment, which prohibits their pharmacological screening using traditional, whole-cell patch clamp electrophysiology and soluble Ca^2+^-indicator dyes^17,27^. Like renal polycystins, PKD2L1 also functions as cilia channel, but it is primarily expressed in hippocampal interneurons and cerebrospinal fluid-contacting neurons in the spinal cord^19,28,29^. Unlike PKD2, the subcellular distribution of PKD2L1 is less restricted— forming channels in the neuronal soma or plasma membrane when heterologously expressed in cell lines^17^. In this report, we leverage this feature and use PKD2L1 as an archetypal polycystin channel to carry out high-throughput screen (HTS) electrophysiology for high-potency chemical matter against these elusive targets. Recent advancements in automated patch-clamp technology have made electrophysiology screening for TRP channel ligands a more viable approach^30^. These systems provide GΩ-seal quality data, comparable to manual electrophysiology, but simultaneously voltage clamp hundreds of cells in planar configuration. Combined with automated liquid handling capabilities, these instruments allow for screening of entire chemical libraries and thorough assessments of a test article’s potency and biophysical mechanism of action against PKD2L1. Our screen results identify four potent PKD2L1 antagonists with divergent chemical core structures from separate drug classes, including local anesthetics and class 1C antiarrhythmics that target voltage-gated sodium channels (Na_v_). In silico analysis of drug docking to PKD2L1 cryo-EM structures pinpoints high-affinity occupancy sites within the vestibule of the pore domain, which are conserved in Na_v_ channels. Our study spotlights the PKD2L1 pore vestibule and lateral fenestrations as promiscuous and state-dependent receptors for chemical antagonism of polycystin channels. Furthermore, it delineates a shared molecular pharmacology between Na_v_ and polycystins in mechanistic and structural terms, which are driven by state-dependent conformational changes, initiated by membrane depolarization.

## RESULTS AND DISCUSSION

### HTS identifies potent PKD2L1 antagonists with diverse chemical core structures

Functional screens of TRP channels are frequently mired by spontaneous sensitization and desensitization phenomena— biological features that can lead to false-positive and false-negative interpretations of drug efficacy^31–33^. Polycystins (PKD2 and PKD2L1) display both calcium-dependent modulation (CDM) and irreversible calcium-dependent desensitization (CDD) when natively expressed in cilia membranes or when overexpressed^34,35^. High-throughput planar electrophysiology systems require low-capacity calcium buffering and the use of fluoride within their intracellular saline to support high-resistance seals in whole-cell configurations^36,37^. When testing HTS saline compositions under manual patch-clamp, transiently expressed PKD2L1 channels recorded from the plasma membrane of HEK cells exhibit rapid CDM and CDD phenomena. These manifest as a transient potentiation of tail currents activated by depolarization trains (+20mV, 0.2Hz) and a subsequent loss (within five minutes) of all tail currents **(Figure S1A, B)**. Previously, a complete alanine scan of the PKD2L1 pore domain identified the pore turret F514A mutant, which abolishes CDD and produces stabilized tail current magnitudes for over 15 minutes under HTS conditions **(Figure S1B)**^38^. Before moving forward with the F514A mutant in our drug screening campaign, we validated its pharmacological sensitivity to previously identified PKD2L1 antagonists (dibucaine, lidocaine, Gd^3+^, and La^3+^)^35^. The resulting dose-response relationships show that against all four antagonists evaluated, WT PKD2L1 and F514A channels display similar potency (1-3x) of current inhibition (IC_50_) **(Figure S1C-F, Table S2)**. Further, we verified the sensitivity of F514A to these same compounds under automated patch clamp (Nanion Synchropatch) and manual patch clamp (Molecular Devices) configurations and observed that for all antagonists (except Gd^3+^) potencies against F514A remained consistent **(Figure S2A-D, Table S2)**. We suspect the reduction of Gd^3+^ potency (IC_50_ = 0.9 μM to 6.5 μM) was caused by precipitation of GdF_3_ formed by electrolysis with the fluoride present in the HTS internal saline, and thus did not include additional trivalent ions in our screening library. Together, these results confirm the suitability of our approach— using the non-desensitizing mutant F514A for electrophysiology-based HTS to identify pharmacological chemical modulators of polycystin channels.

To assemble our polycystin screening library (100 compounds total), we combined small molecules from commercially available channel-ligand screening collections (SCREEN-WELL®, Enzo Life Sciences) with individually sourced prototypic or clinically used drugs (Millipore Sigma) with established channel pharmacology **(Table S1)**. To identify ligands with reproducible and high potency efficacy against PKD2L1, we assessed the effects of each test compound at 1 μM concentration against 6-39 separate cells over several cell line passages. From this first pass trial, we identified a chemically divergent set of compounds (U-50488, propafenone, tetrandrine, and AM-92016) which significantly inhibited polycystin tail currents **(Figure 1A)**. Additional concentration-response experiments defined their potency of inhibition (IC_50_) ranging from 1.0-3.8 μM **(Figure 1B, Figure S1C, E)**. Notably none of the screened compounds enhanced PKD2L1’s voltage-activated tail currents and less than 4% had antagonist properties, suggesting that as a drug target, the channel is not prone to indiscriminate chemical activation or inhibition. Among the identified PKD2L1 antagonists, dibucaine, propafenone and tetrandrine are canonical voltage-gated sodium channel (Na_v_) and calcium channel (Ca_V_) antagonists with antiarrhythmic and local anesthetic properties and established receptor sites within their molecular targets^39–43^.

**Figure 1.**
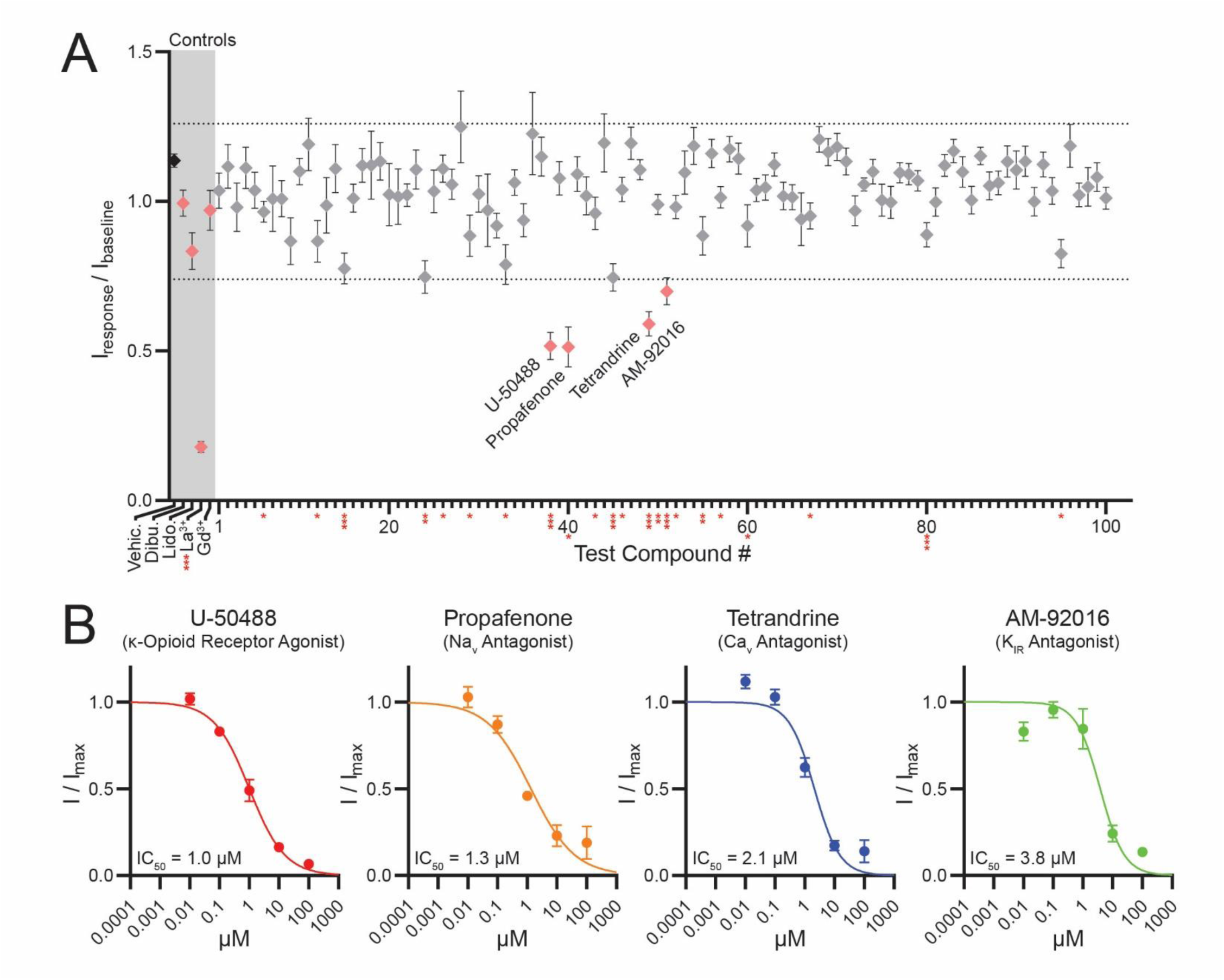
High-throughput drug screen identifies four potent PKD2L1 antagonists. (**A**) Mean post-treatment F514A tail currents, normalized to pre-treatment amplitude. Asterisks denote compounds with a statistically significant change in amplitude relative to currents from vehicle treated, cells (*p < 0.05, **p < 0.005, ***p < 0.0005, unpaired t-test). Dotted lines denote a ± 25% shift from baseline current. Grey overlay distinguishes control and previously identified PKD2L1 antagonists. (controls DMSO, La3+ n = 7-109 cells; test compounds n = 6-39 cells, Error = SEM) (**B**) Drug concentration-response relationships for F514A when treated with screen-identified antagonists from **A**. Data were fit to the Hill equation to calculate IC_50_ values. (n = 6-17 cells, Error = SEM)

### In silico structural analysis predicts antagonist receptor sites within the pore vestibule

Given the chemical diversity of our identified potential ligands and the paucity of structural information on potential receptor sites within the PKD2L1 channel, we used previously determined cryo-EM structures to perform in-silico docking analyses for all four hits using AutoDock4.2^44^. Blind docking results exhibited a propensity toward stable binding within the PKD2L1 pore domain (data not shown). With this in mind, we performed targeted docking of the pore domain and generated clustering histograms for the resulting conformations **(Figure 2A)**. U-50488 and tetrandrine docking produced conformational clusters with a narrow range of predicted binding energies and averages of −7.16 kcal/mol and −8.65 kcal/mol, respectively **(Figure 2A-B)**. In contrast, propafenone and AM-92016 analyses generated docking conformations with greater variance in binding energies, and averages of −4.65 kcal/mol and −6.65 kcal/mol, respectively **(Figure 2A-B)**. Next, we visualized the 10 highest ranked conformations for each compound to better interpret their proposed binding sites. These models predict that for U-50488 and propafenone, stable binding conformations cluster within the upper surface of the pore vestibule **(Figure S3A-B)**. For tetrandrine, the highest ranked conformations form a cluster within the bottom of the pore vestibule **(Figure S3C)**. AM-92016 shows the least consistent grouping, with predicted binding conformations distributed throughout the pore vestibule **(Figure S3D)**. To gauge the relevance of these docking conformations to our in vitro data, we compared AutoDock-predicted K_i_ values to our in vitro IC_50_ values. Predicted K_i_’s for the most stable conformations of U-50488 (K_i_ = 0.9 μM), tetrandrine (K_i_ = 0.3 μM), and AM-92016 (K_i_ = 1.5 μM) either fell in agreement with or predicted a greater potency than measured in vitro (IC_50_ = 1 μM, 2.1 μM, and 3.8 μM, respectively) **(Figure 1B, Table S2, Table S3)**. In contrast, the predicted K_i_ of propafenone (K_i_ = 11.9 μM) showed a 10-fold lower potency than observed in vitro (IC_50_ = 1 μM) **(Figure 1B, Table S2, Table S3)**. These results are consistent with the known limitations of the software when docking molecules with high torsion numbers, and suggest that for three of our four compounds, the predicted dynamics of their most stable docking conformations agree with our in vitro observations. Notably, for all compounds except tetrandrine, the top-ranked docking conformations localized to the upper pore domain, implying the existence of a binding site with favorable interactions for multiple drug classes **(Figure 2B)**. Confident that the top-ranked conformations predicted binding activity comparable to that observed in vitro, we analyzed our docked structures with the Protein-Ligand Interaction Profiler (PLIP)^45,46^. Across all four drug-bound models, hydrophobic interactions provided the bulk of stabilizing forces **(Figure 2C)**. Pore vestibule residues F556 and F549 participate in pi-stacking interactions with tetrandrine’s aromatic rings, while the backbone oxygen molecules of I520 and L552 form hydrogen bonds with propafenone and AM-92016. Interestingly, the aromatic rings of top-ranked conformations for U-50488, propafenone, and AM-92016 are coordinated by sidechain phenyls of F549 and F524. The proximity to these residues suggests the potential to swap to alternate conformations with the ability to pi-stack. Together, the modeling analyses suggest the pore vestibule is a shared receptor site for a diverse set of chemical antagonists of PKD2L1 channels.

**Figure 2.**
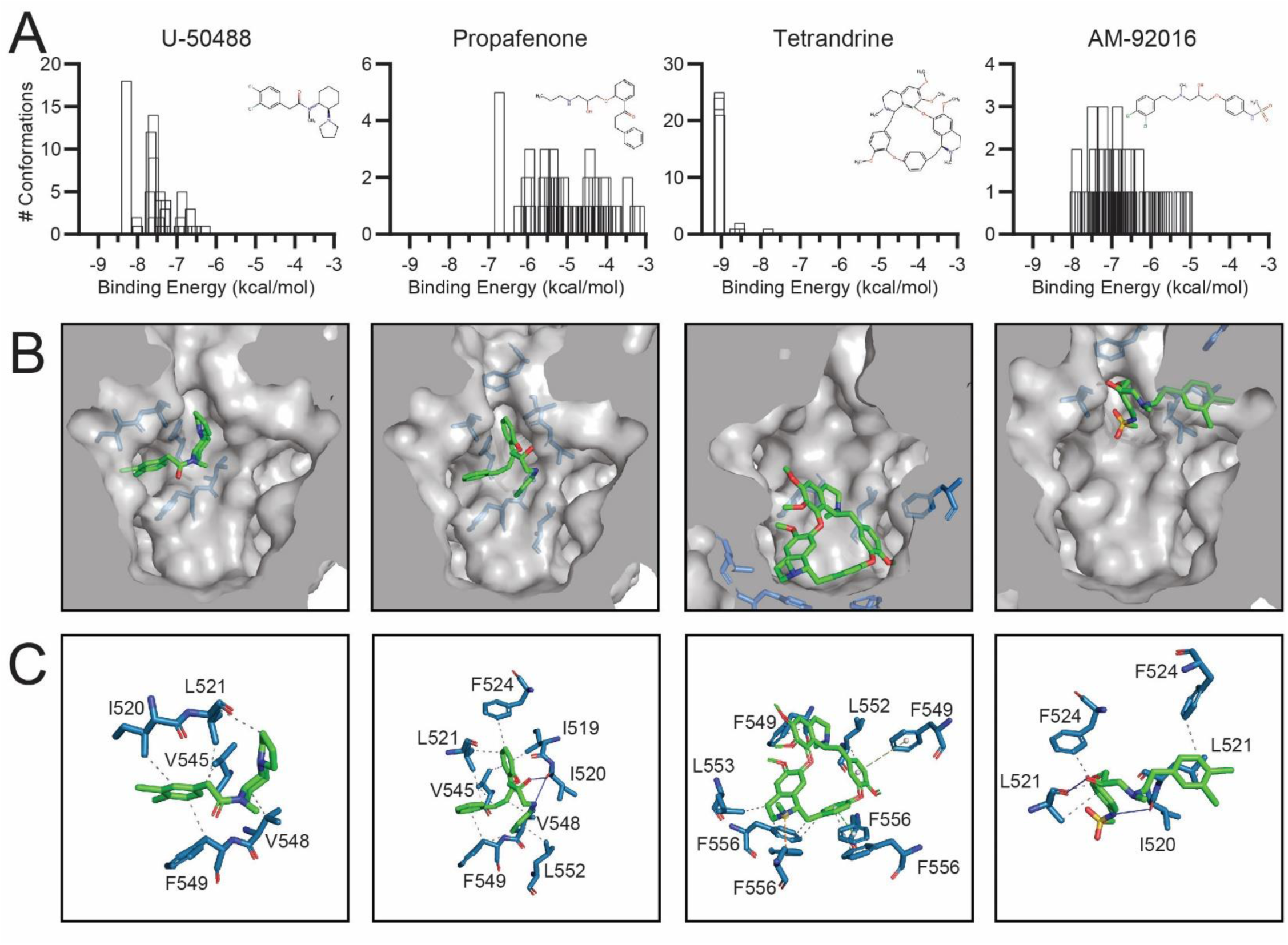
In silico docking predicts drug binding within the PKD2L1 pore vestibule. (**A**) Clustering histograms of screen hits. Histograms represent the results from 100 runs of docking analysis against PKD2L1 (PDB: 6DU8), grouped by conformational clusters and ranked according to the lowest binding energy in each cluster. Insets present the chemical structure for each screen hit^10^. (**B**) Surface representations of highest-ranked docking conformation for each screen hit. (**D**) Stick representations of molecular interactions between screen hits and binding site residues, analyzed via PLIP.

### PKD2L1 lateral fenestration residues forms a local anesthetic-antiarrhythmic receptor site

As demonstrated from our pharmacological screen, PKD2L1 currents are sensitive to sodium channel antagonists (dibucaine, lidocaine, propafenone)— drugs that are clinically used as local anesthetics (LAs) and/or class 1C antiarrhythmics (AAs). The molecular location of their binding site was first described in mutagenesis studies, but subsequent structural determinations in prokaryotic Na_v_ orthologs (e.g. Na_v_Ab) have defined the upper pore vestibule as the receptor site responsible for LA-AA drug action^47,48^. Here, these molecules interact with residues lining the lateral fenestrations within the Na_v_ pore domain, formed by S5-S6 transmembrane helical segments. The pore domains of homotetrameric PKD2L1 and Na_v_ channels are structurally homologous (32% by residue identity and 58% by character), and both channel types have lateral fenestrations with comparable dimensions in their high-resolution structures **(Figure 3A)**^49^. Within the lateral fenestrations, several key Na_v_ LA-AA receptor site residues are conserved in PKD2L1 channels and retain their vestibule-facing structural orientation, as well as the general hydrophobicity of their neighboring residues **(Figure 3B-C)**. To predict if a Na_v_-like dibucaine binding site is indeed shared by PKD2L1, we performed in silico docking analysis and observed enhanced affinity for the closed state channel structure (K_i_ = 5.9 μM) compared to open state (Ki = 10.5 μM) at overlapping receptors sites within the upper pore vestibule, near the lateral fenestrations **(Figure 4B)**. Similar to results from our screen hits, PLIP analysis predicted that binding of dibucaine within PKD2L1 relies primarily on hydrophobic interactions between the ligand and upper pore residues **(Figure 4B)**. Additionally, residue F549 is predicted to participate in both hydrogen bonding and pi-stacking with dibucaine **(Figure 4B)**. Considering these docking results, we asked whether mutagenesis of this site could alter drug antagonism in functioning PKD2L1 channels. A previous mutagenic screening of the PKD2L1 pore domain reported that F549A, V, and S mutations all result in the complete loss of channel function— therefore, these could not be tested in our electrophysiology assays^38^. Guided by this knowledge, we chose to target less disruptive substitutions, including hydrophobic residues with comparable sidechain lengths (F549I, L) and aromatic residues that maintained the potential for pi-stacking (F549H, W). Of these, the I, L, and H substitutions all produced functional currents when expressed in mammalian cells **(Figure 4C)**. When comparing the dibucaine potency of current inhibition, F549L (IC_50_ = 10.5 μM) produced little to no change compared to wildtype (IC_50_ = 8 μM), whereas H (IC_50_ = 17.5 μM) and I (IC_50_ = 68.9 μM) substitutions showed a 2 to 10-fold reduction in potency **(Figure 4D)**. Using the change in IC_50_ as an apparent change in K_i_, we calculated a loss of binding affinity (ΔG) of 5.3 kcal/mol for the isoleucine mutation **(Figure 4E, Table S4)**. Due to the overlapping docking conformations for U-50488 and dibucaine, we evaluated U-50488 inhibition of F549I and observed a 2x loss in potency, relative to wildtype PKD2L1 **(Figure S4A-B)**. Together, these results support the conclusion that PKD2L1 and Na_v_s share conserved lateral fenestrations in the pore domains, which form receptor sites for local anesthetics, antiarrhythmics and other chemical antagonists, such as U-50488.

**Figure 3.**
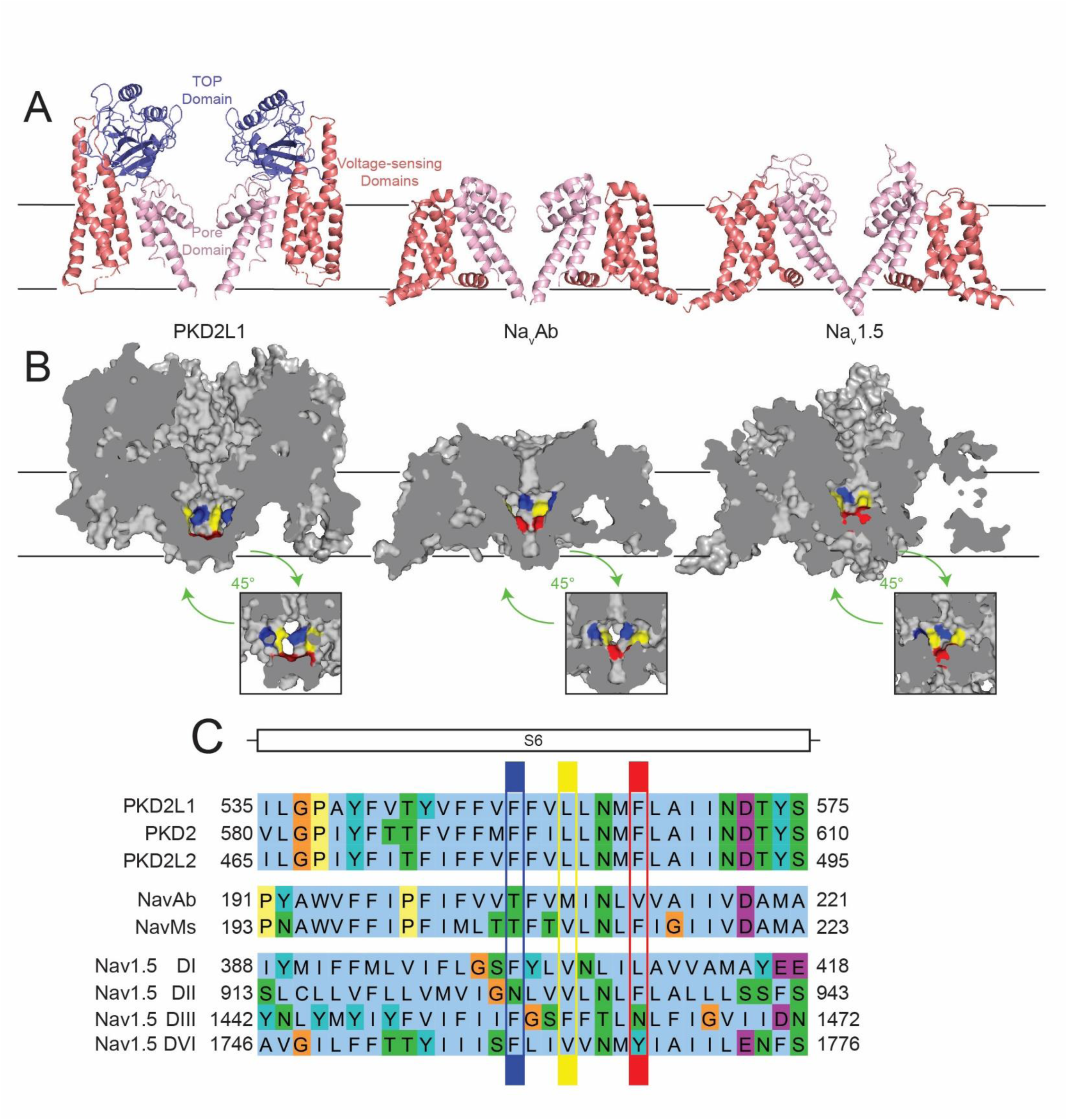
Structural homology between PKD2L1 and Na_v_ channels implies shared drug binding properties. (**A**) Cartoon representations of two isolated domains from PKD2L1, Na_v_Ab, and Na_v_1.5 high-resolution structures, with channel regions colored to highlight the pore domain (pink), voltage-sensing domains (red), and TOP domain (blue)^10,59,60^. (**B**) Cross section-view of PKD2L1, Na_v_Ab, and Na_v_1.5 surface representations. Insets provide alternate viewing angle of the pore domain’s interior surface. Colors denote locations Na_v_Ab drug-binding residues: T206 *blue*, M209 *yellow*, V213 *red*. (**C**) Sequence alignment of the S6 transmembrane segments of PKD2L1, PKD2, Na_v_Ab, Na_v_Ms, and Na_v_1.5. Sequence residues colored according to ClustalX scheme, with highlights corresponding to colored residues in **B**.^61^

**Figure 4.**
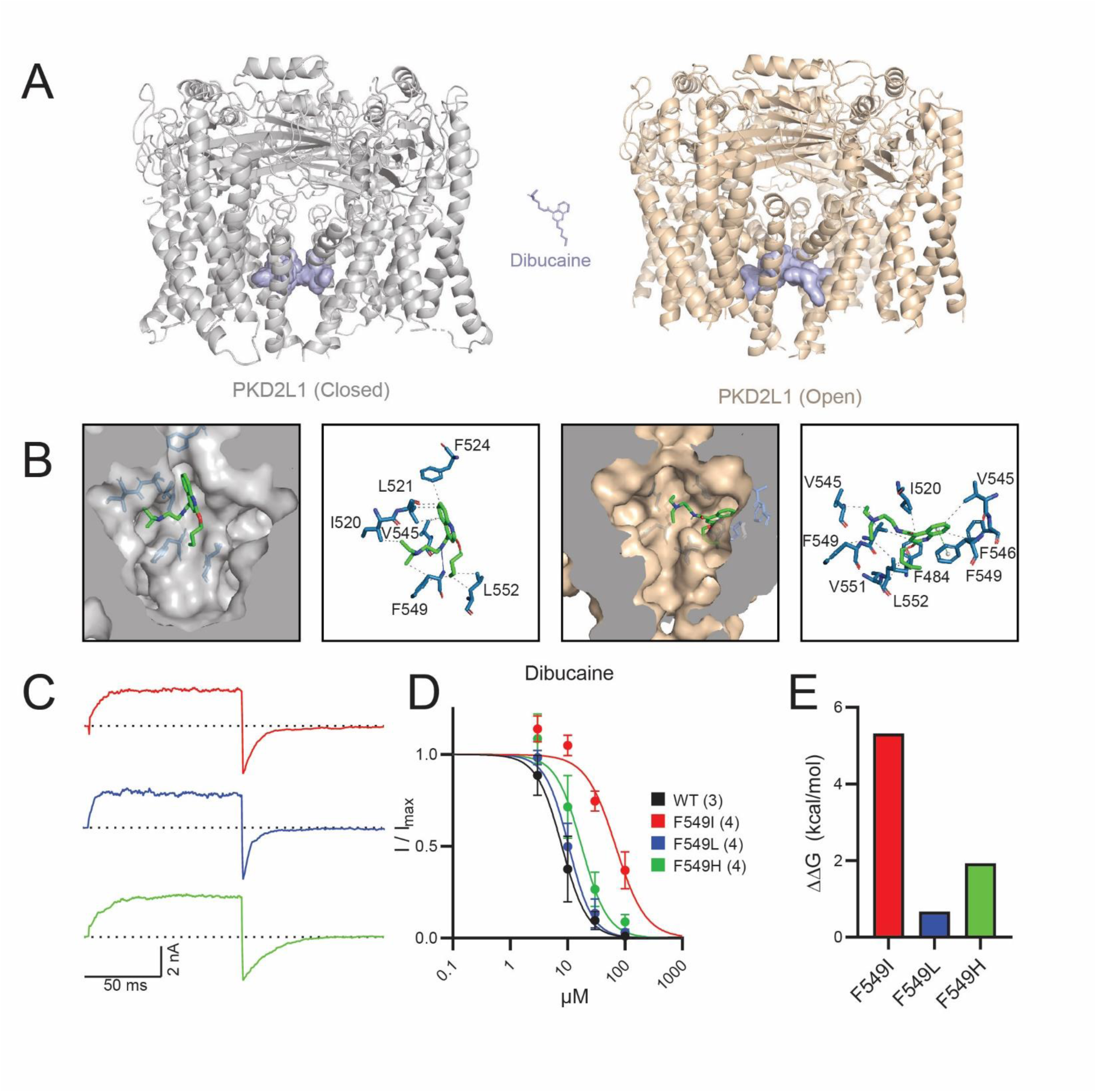
PKD2L1 contains a Na_v_-like local dibucaine binding site. (**A**) Cartoon representations of PKD2L1 in closed (PDB: 6DU8) and open (PDB: 5Z1W) conformations^9,10^. Blue surfaces highlight the ten highest ranked docking positions obtained for dibucaine in Autodock4.2. (**B**) Surface representations of the highest-ranked binding positions for dibucaine. Stick representations of proposed molecular interactions between dibucaine and binding site residues, analyzed with PLIP. (**C**) Sample current traces from functional F549 mutations. (**D**) Dose-response curves for dibucaine inhibition of PKD2L1 WT and F549 mutant currents. (n = 4-6 cells, Error = SEM). (**E**) Change in dibucaine binding affinity for functional F549 mutants, calculated from the change in IC_50_ values relative to WT PKD2L1.

### State-dependent mechanism of PKD2L1 channel inhibition

Na_v_ antagonism by LAs and AAs is characterized by frequency-dependent (i.e., use-dependent) inhibition, a feature that underlies their clinical efficacy and clinical safety^47,50^. Na_v_s under high-frequency depolarizations show enhanced LA-AA receptor potency and accelerated kinetics of inhibition, which suggests greater ligand access to the lateral fenestration receptor site in the open or inactivated states. These characteristics allow for selective inhibition of aberrantly hyperexcitable neurons or cardiomyocytes, without impairing the rising phase of action potentials from unaffected cells^50^. To determine if local anesthetic access to the PKD2L1 lateral fenestration receptor is also state-dependent, we applied 100 ms depolarizations (100 mV) at different frequencies in presence of 10 µM dibucaine **(Figure 5A)**. The rate of inhibition was three-fold faster at 2 Hz (τ = 8 s) than at 0.2 Hz (τ = 25.8 s) stimulus, consistent with frequency-dependent inhibition and greater drug accessibility to the lateral fenestration receptor in the open state. Examination of the cryo-EM structures of WT PKD2L1 and modeled F549 mutants indicate that the lateral fenestration becomes sealed in the open state, suggesting small molecules become trapped when the channel transitions out of the closed state (**Figure S5A-B**). Once bound to the lateral fenestration receptor, we hypothesized two molecular mechanisms which could result in channel inhibition. LAs (or other analogues) could either bind and directly plug the ion conducting pathway of the channel or bind and stabilize PKD2L1’s non-conducting inactivated state. When examining our models of dibucaine bound to the fenestration receptor (open state), orientation of the drug molecules did not encroach into central vestibule, suggesting that the mechanism is not blockade of the incoming cations. To assess the alternative hypothesis, we compared PKD2L1 voltage-dependent inactivation in control and dibucaine-treated conditions using a 2 second variable prepulse (−60 to 100 mV), followed by a test pulse (100 mV) to open the available channels **(Figure 5B)**. Here, we observed a significant hyperpolarizing shift (−69 mV) in voltage-dependent inactivation after dibucaine treatment (Inactivation V_1/2_, control = 49 mV, dibucaine = −20 mV). Taken together, these results demonstrate LA access the PKD2L1 lateral fenestration receptor site in the open-inactivated states. Once they occupy the lateral fenestration receptor site, LAs stabilize the non-conducting inactivated state of the channel, which causes inhibition of the current (**Figure 6A**).

**Figure 5.**
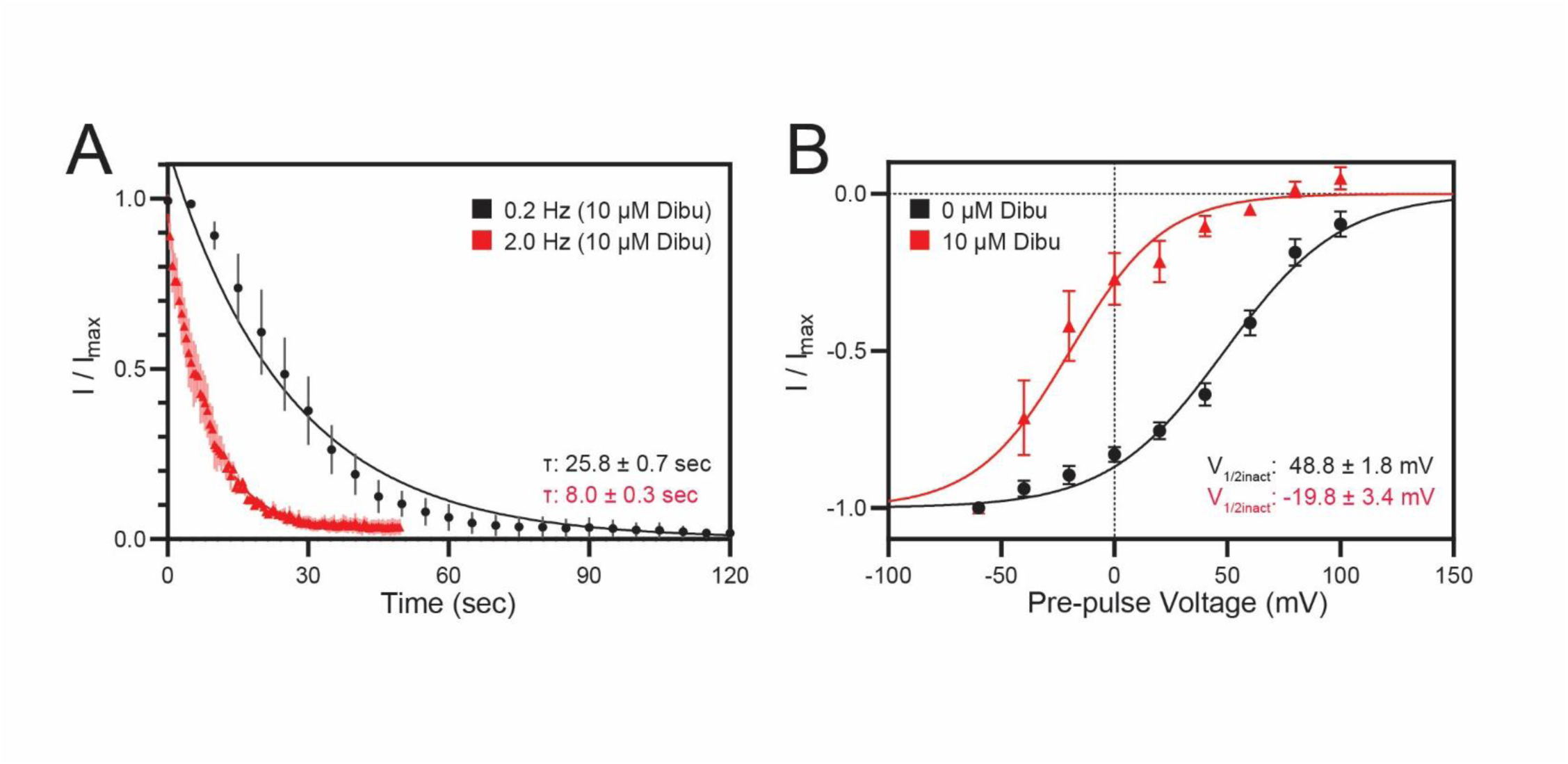
Dibucaine shows use and voltage-dependent inhibition of PKD2L1. (**A**) Time course of PKD2L1 inhibition by 10 µM dibucaine at 0.2 and 2 Hz sweep frequencies. Time constants (tau) were calculated by fitting tail currents to a one-phase model of exponential decay. (n = 4 cells, Error = SEM) (**B**) Normalized current-voltage relationship for PKD2L1 when treated with 0 and 10 µM dibucaine. V_1/2_ for each condition was estimated by fitting data to a two-state Boltzmann function. (n = 5 cells, Error = SEM)

**Figure 6.**
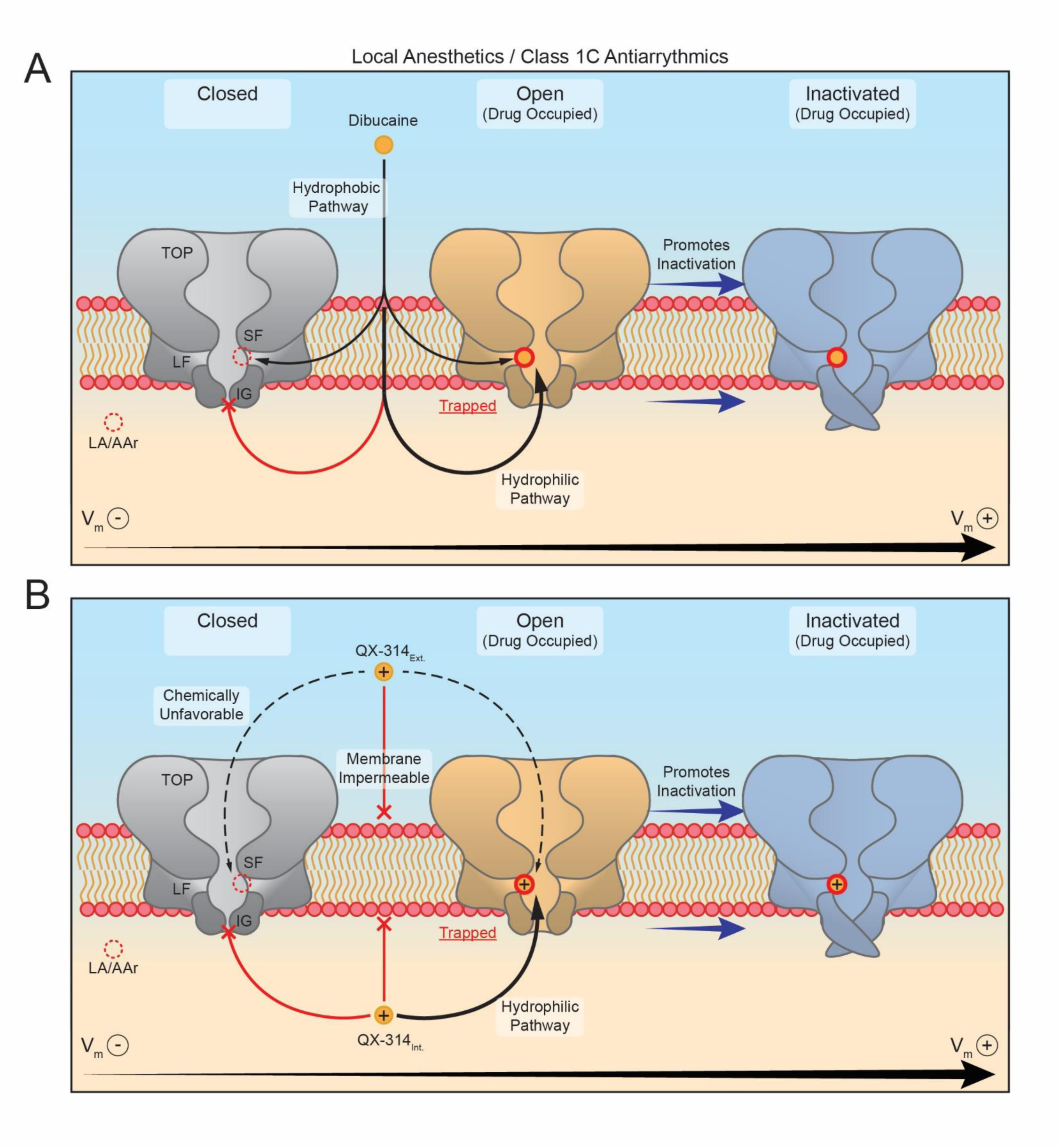
A hypothetical model of drug access to the PKD2L1 lateral fenestration binding site. (**A**) LA-AA receptor access to the lateral fenestration binding site preferentially occurs via the inner gate (IG), as evidenced by enhanced inhibition of PKD2L1 during high-frequency opening events. For drugs with high lipid solubility, the lateral fenestrations (LF) present an additional hydrophilic access path to the LF receptor, as in Na_v_s. (**B**) Membrane-impermeable local anesthetics (QX-314) show bilateral access to the local anesthetic binding site, with intracellular treatment producing near instantaneous inhibition, while extracellular treatment produces exceedingly slow inhibition. Due to QX-314’s inability to traverse through the cell membrane, this extracellular inhibition indicates a path through the tetragonal opening for polycystins (TOP) and selectivity filter (SF) to access the LF receptor. Due to the delayed inhibition produced by extracellular QX-314, this SF pathway is likely much less chemically favorable than the intracellular hydrophilic pathway.

Dibucaine and the other screen-identified antagonists have high octanol-water partition coefficients (logP > 3) and can permeate cell membranes. Since the lateral fenestration is located within the transmembrane vestibule, we propose that drug molecules access this receptor site either by traversing within the cell membrane and through the lateral fenestration (hydrophobic pathway) or by passing through membrane to access the open pore from the cytosolic side (hydrophilic pathway) (**Figure 6B**). To test drug access to the PKD2L1 lateral fenestration receptor, we explored PKD2L1 current inhibition by membrane impermeable QX-314 (a lidocaine derivative), previously used to define the mode of Na_v_ inhibition by local anesthetics^47,51^. When introduced to the internal side of channel via the patch electrode, QX-314 produced an immediate and total inhibition of PKD2L1 currents, reminiscent of results from classical experiments characterizing Na_v_ channel inhibition (**Figure S6A**). To determine if QX-314 could access the lateral fenestration receptor by passing through the inner gate, we measured the pore radii of PKD2L1 closed and open structures using the HOLE program (**Figure S6B, C**)^52,10,11^. Here, the inner gate radius is restricted to 0.55 Å in the closed state and dilates to 2.6 Å upon channel opening, which is wide enough to pass molecules from the cytosolic side and may explain the observed open state-dependence of local anesthetic inhibition. When applied to the external side of the channel via the bath solution, QX-314’s effect on PKD2L1 currents were limited, requiring > 7 minutes to reach steady state inhibition (**Figure S6A**). These results suggest drug passage through the selectivity filter from the external side is less chemically favorable and cannot confer state-dependent inhibition based on its less dynamic and wide aperture (R= 1.95-2.6 Å). Taken together, we propose that uncharged local anesthetics and class 1C antiarrhythmics can access the lateral fenestration receptor site through hydrophobic or hydrophilic pathways— either by traversing through the membrane fraction or maneuvering into the pore vestibule when the intracellular gate opens.

## SUMMARY

Using high throughput electrophysiology, we identified a chemically diverse group of potent PKD2L1 antagonists, including several with known voltage-gated ion channel pharmacology. In-silico docking analysis and functional mutagenesis identified fenestrations within the PKD2L1 pore domain as a binding site for these compounds. Previous studies of TRP channel pharmacology have highlighted unique ligand binding pockets within their voltage-sensor (or voltage-sensor like) and cytosolic domains, but receptors within the cavities of the pore vestibule have not been previously reported^53–55^. Polycystins are the earliest-evolved subfamily of TRP channels, with orthologs present in the genomes of bacteria, yeast, and protists. These channels diverged from voltage-gated channels ≈750 million years ago^56,57^. Based on their common evolutionary origins, it is reasonable to expect that along with several conserved structural features, that polycystins would also share pharmacological sensitivities with Na_v_ and Ca_v_ channels^58^. It is important to note that lateral fenestrations and other structural features which confer voltage-dependent gating (e.g., gating charges) found in the polycystin subclass are not present in other subfamilies of TRP channels. Thus, these features are unique to polycystins and can be leveraged to develop selective agents that state-dependently target members of this TRP subclass. Importantly, the PKD2L1 lateral fenestration site (e.g., F549) is also conserved in PKD2— whose germline variants are associated with 15-20% of patients with ADPKD. Notably, we did not identify any polycystin channel activators despite screening a diverse chemical library. Given that ADPKD is considered a loss of function disease, the therapeutic value for the identified polycystin antagonists and their receptor sites is uncertain. Nonetheless, our study clearly establishes the utility of the HTS electrophysiology approach and the opportunity to identify prototypic polycystic drug activators by expanding the drug screening library.

## METHODS

### Vector Expression and Mutagenesis

18-24 hours prior to manual recordings, polycystin channel cDNA was co-transfected alongside enhanced green fluorescence protein (EGFP) cDNA using Lipofectamine 2000 (Invitrogen). To generate stable cells for automated recordings, HEK 293 cells were infected with lentivirus expressing PKD2L1 F514A and IRES GFP under a CMV promoter (Vector Builder). After 48 hours, the resulting stable cells were selected with puromycin (Thermo Fisher Scientific) and maintained in Dulbecco’s modified Eagle’s medium (GIBCO) supplemented with 10% fetal bovine serum (Sigma-Aldrich) and 50 mg/mL streptomycin (Cytiva). Plasmids for PKD2L1 mutations were generated via site-directed mutagenesis of a WT PKD2L1 plasmid using the Q5 Site-Directed Mutagenesis Kit (New England Biolabs). Mutagenic primers were designed through the Q5 designer tool (New England Biolabs) and are listed in **Table S5**. For all engineered amino acid substitutions, successful mutagenesis was verified with whole-plasmid sequencing (Plasmidsaurus).

### Chemical library

High-throughput electrophysiology screen was performed using the 70 compounds included in the Screen-Well Ion Channel Ligand Library (BML-2805-0100, Enzo) and 30 individually sourced compounds. All compounds in the Enzo library had stock concentrations of 10mM in either DMSO or water, while individually sourced compounds were reconstituted to either 1 mM or 10 mM concentrations in DMSO or water. Aliquots of each stock solution were prepared prior to screen experiments to avoid contamination and freeze-thaw degradation of original stocks. When not in use, stock solutions were stored at −80 °C and aliquots were stored at −20 °C. Prior to each run of HTS, test article working solutions were prepared fresh in assay buffer (extracellular recording solution) to achieve a final concentration of 1 µM in recording wells. For concentration-response experiments, test articles were first diluted at log/half-log increments in DMSO/water to produce a series of five test concentrations. To ensure validity of screen-identified hits, subsequent concentration-response experiments were performed with externally sourced samples. All compounds and sources can be found in Table S1.

### Manual Patch Clamp

Whole-cell patch clamp recordings were performed with an Axiopatch 200B amplifier, Digidata 1550B digitizer, and pClamp 11 recording software (all Molecular Devices). Borosilicate glass capillaries (1B150F-4, World Precision Instruments) were pulled one a P-1000 pipette puller (Sutter Instruments) to make recording electrodes with 2-5 MΩ tip resistances. Extracellular recording solution contained: 150 mM NaCl, 10 mM HEPES, and 2 mM CaCl_2_. Pipette solutions contained either: 80 mM CsMES, 20 mM NaCl, 15 mM BAPTA, 10 mM HEPES, 5 mM EGTA, and 2 mM MgCl_2_ for WT PKD2L1 and F549 mutant cells (except where otherwise noted) or: 110 mM CsF, 10 mM CsCl, 10 mM CsCl, 10mM HEPES, and 20mM EGTA for all other experiments. All solutions were adjusted to a final pH of 7.4 and an osmolarity of 300 mOsm (± 5mOsm) with mannitol. All experiments were performed at 21 °C. Tail currents were elicited through a voltage-step protocol with a holding potential of −60 mV, a 100ms step to +100 mV, and a repolarization to −60 mV for wildtype PKD2L1. For PKD2L1 F514A, this was adjusted to a −80 mV holding potential and 120mV step to account for the altered voltage-dependence of opening, as previouly reported^38^. To minimize voltage error, cells were excluded from analyses if I_leak_ exceeded −200 pA at V_holding_, or if R_series_ exceeded 10 MΩ. Normalized current inhibition was calculated from the ratio of pre-treatment current amplitudes and post-treatment steady-state current amplitudes, expressed as I/I_max_. Concentration-response plots were fit to the Hill equation (Y = Base + (Max-Base)/(1+(IC_50_/x)^HillSlope)) to evaluate the potency (IC_50_) of inhibitory compounds. Voltage-dependence of inactivation was analyzed through fitting a two-state Boltzmann (I/I_Max_=1/(1+exp((V_1/2_-V)/k)) distribution to normalized tail current data generated by a pre-pulse protocol consisting of a 2-second variable prepulse (−60 to 100mV), followed by a test pulse (100mV).

### Automated Patch Clamp

Whole-cell planar patch recordings were performed on a Syncropatch 384I (Nanion Technologies) automated patch clamp platform using S-type 384 well 1X chips with 2-4 MΩ resistances (Nanion Technologies). Pulse protocols and data collection were controlled via PatchController384 v3.2.0.27 (Nanion Technologies). Extracellular recording solution contained: 140 mM NaCl, 10 mM HEPES, 5 mM glucose, 4 mM KCl, 2 mM CaCl_2_, and 1mM MgCl_2_ with a final pH of 7.4, adjusted with NaOH. Seal enhancer solution contained: 80 mM NaCl, 35 mM CaCl_2_, 10 mM MgCl_2_, 10 mM HEPES, and 3 mM KCl with a final pH of 7.4, adjusted with NaOH. Intracellular solution contained: 110 mM CsF, 10 mM CsCl, 10 mM CsCl, 10 mM HEPES, and 20 mM EGTA, with a final pH of 7.2, adjusted with CsOH. All experiments were performed at 21 °C. Rigorous criteria were used to filter out poor-quality recordings prior to final data analysis. Cells qualified for final analysis if they met the following standards: Seal resistance ≥ 200 MΩ. Series resistance ≤ 20 MΩ. Baseline current between −100 and 100 pA. Tail peak amplitude between −200 pA and −10 nA. During standard planar patch experiments, 384 well chips were partitioned into 12 two-column sections, 10 of which contained test compounds, 1 of which contained a known blocker (LaCl), and one of which served as a DMSO control, for a total of 32 potential replicates per condition. Compounds were identified as hits if 1 µM treatment resulted in 1) a change from baseline to response amplitude of greater than 25% and 2) a statistically significant difference between the mean response of treated and control cells.

### 3D Modeling and In-silico Docking

All in-silico docking models were generated with AutoDock4 v4.2.6^44^. PKD2L1 open (PDB: 6DU8) and closed (PBD: 5Z1W) structures were prepared for docking analysis through removal of water molecules, the addition of polar hydrogens, and the addition of Kollman charges. Atoms were then assigned an AutoDock 4 type. Ligand molecules were optimized in Avogadro, and all rotatable bonds were unlocked prior to docking. All molecules first underwent blind docking, during which the bounding box encompassed the entirety of the channel structure. Based on the results of blind docking, the bounding box in subsequent docking analyses was restricted to regions with the most stable docking conformations. Final docking results were obtained with 100 runs of the Lamarckian Genetic Algorithm under the following parameters: Population Size = 300, Number of Generations = 27000, and Number of Evaluations = 2.5-50 million depending on the torsion # for a given ligand.

### Data Analysis

Data were analyzed via a combination of DataControl384 (Nanion Technologies), pClamp 11 (Molecular Devices), Prism 8 (Graphpad), and Rstudio (Posit). Access to R-scripts used to process data is available upon request.

### Statistical Analysis

Statistical methods used to determine significance are described in corresponding figure legends and reported as *P* values. Briefly, electrophysiology and imaging datasets were analyzed (GraphPad or Origen) using one way ANOVA, Student’s t-tests, two-tailed, paired (equal sample sizes) or unpaired (unequal sample sizes).

### Materials availability statement

All cell expression constructs used in this study are available without restriction upon written request to the corresponding author.

## Data availability statement

Data reported in this paper is deposited and will be made available after publication, without restriction through the NU library ARCH (https://doi.org/10.21985/n2-f3zk-rq43).

---

**Figure S1.**
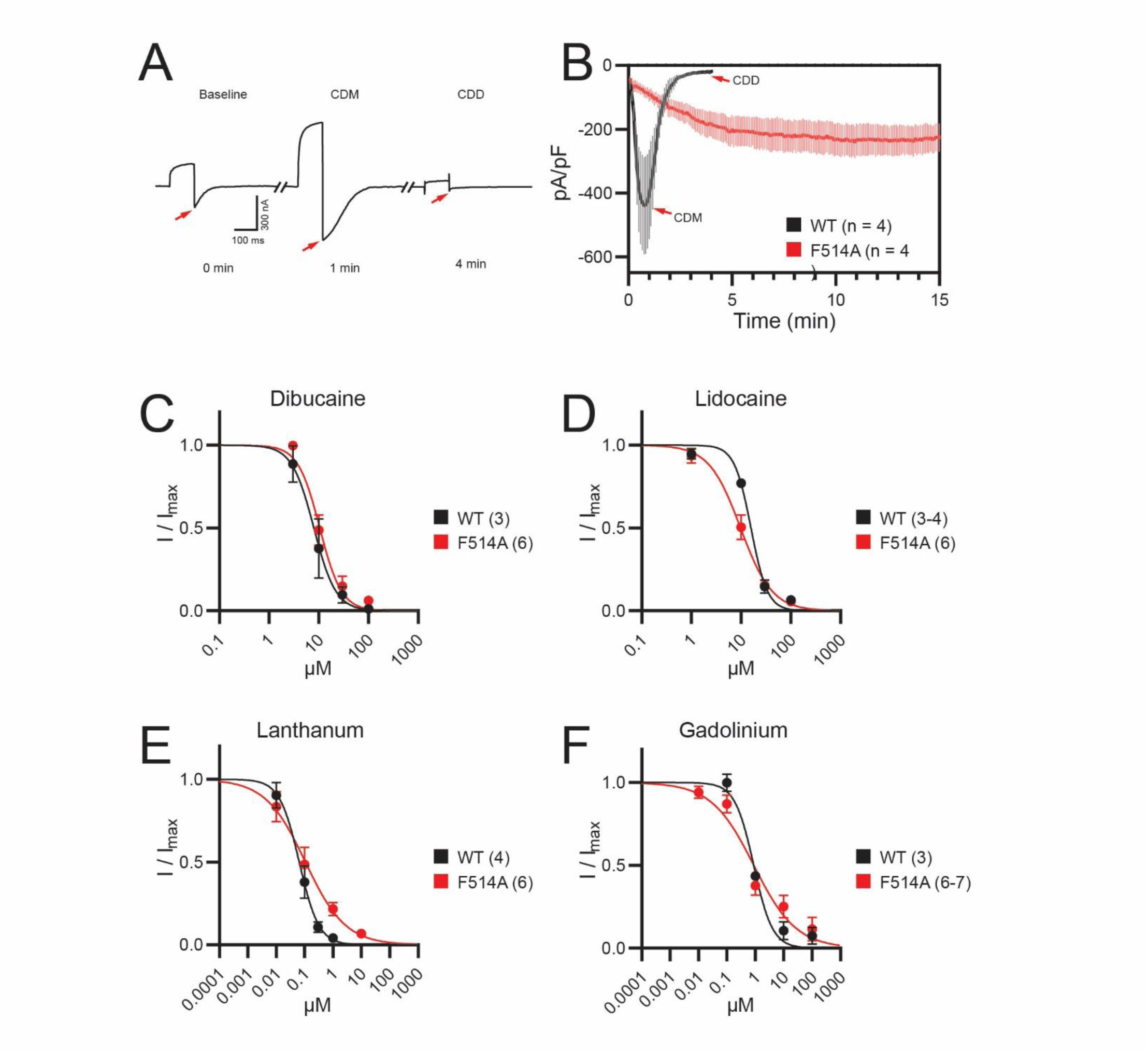
Non-desensitizing F514A mutation maintains drug-binding properties comparable to WT PKD2L1. (**A**) Exemplar current recordings from WT PKD2L1 in 15mM EGTA internal solution. Currents were elicited by 100 ms steps to 100 mV from a holding potential of −80 mV. (**B**) Time course of tail current density over time for WT PKD2L1 and F514A (n = 4 cells, Error = SEM). (**C-D**) Concentration response relationships for inhibition of PKD2L1 and F514A by known PKD2L1 antagonists (n = 3-7 cells, Error = SEM).

**Figure S2.**
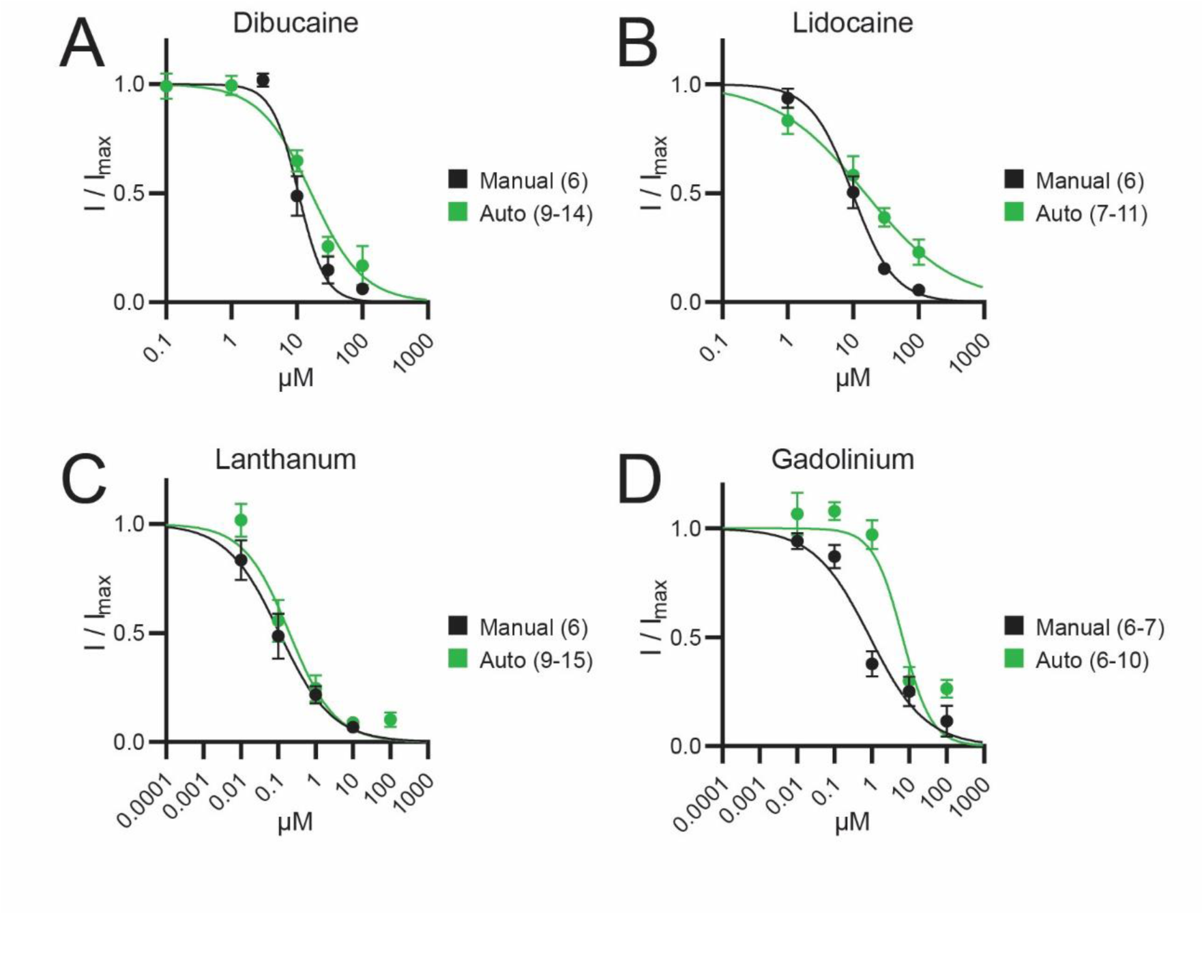
Automated planar patch produces drug responses comparable to manual patch clamp. (**A-D**) Concentration response relationships for inhibition of F514A under manual and automated patch-clamp. (n = 5-15 cells, Error = SEM)

**Figure S3.**
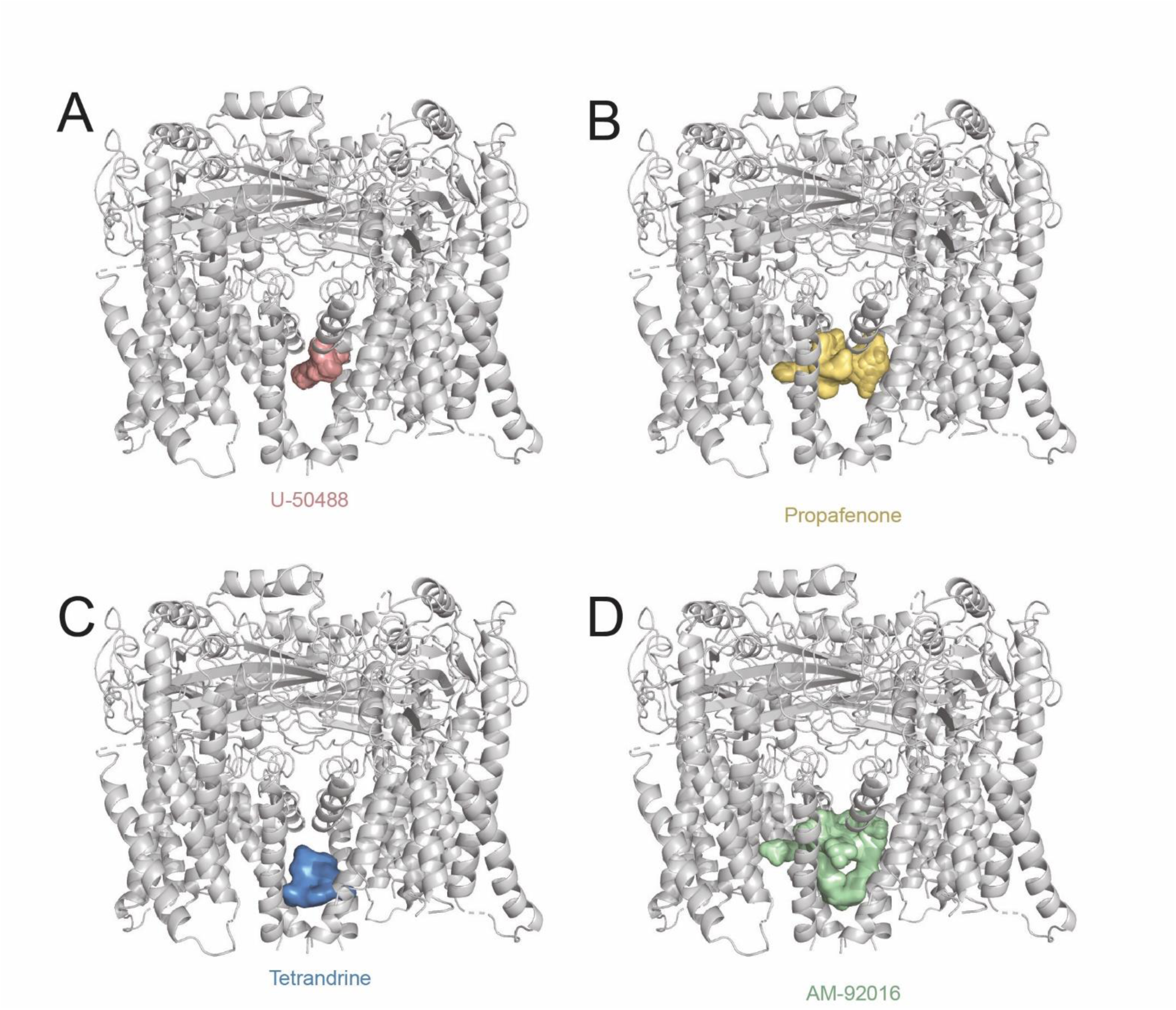
High-rank docking conformations localize to the pore domain for all screen hits. (**A-D**) Cartoon representations of PKD2L1 (PDB: 6DU8) drug-bound models^10^. Colored surfaces highlight the ten highest-ranked docking positions obtained for each PKD2L1 antagonist via AutoDock4.2.

**Figure S4.**
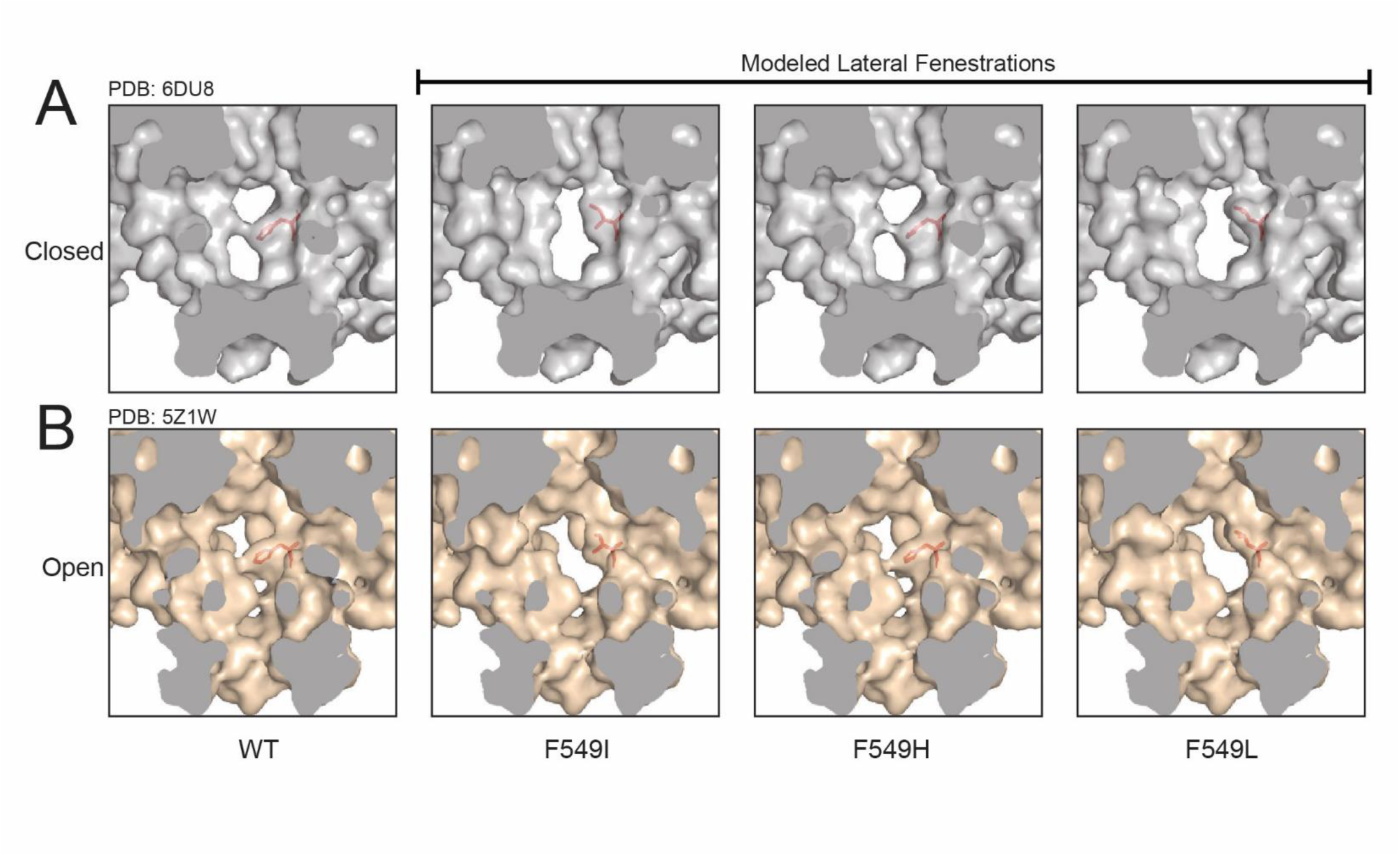
F549 mutations alter the profile of the PKD2L1 window fenestrations. (**A-B**) Cross-section view of the PKD2L1 lateral fenestration site in WT channel structures and F549 variants generated with the Pymol Mutagenesis Wizard.

**Figure S5.**
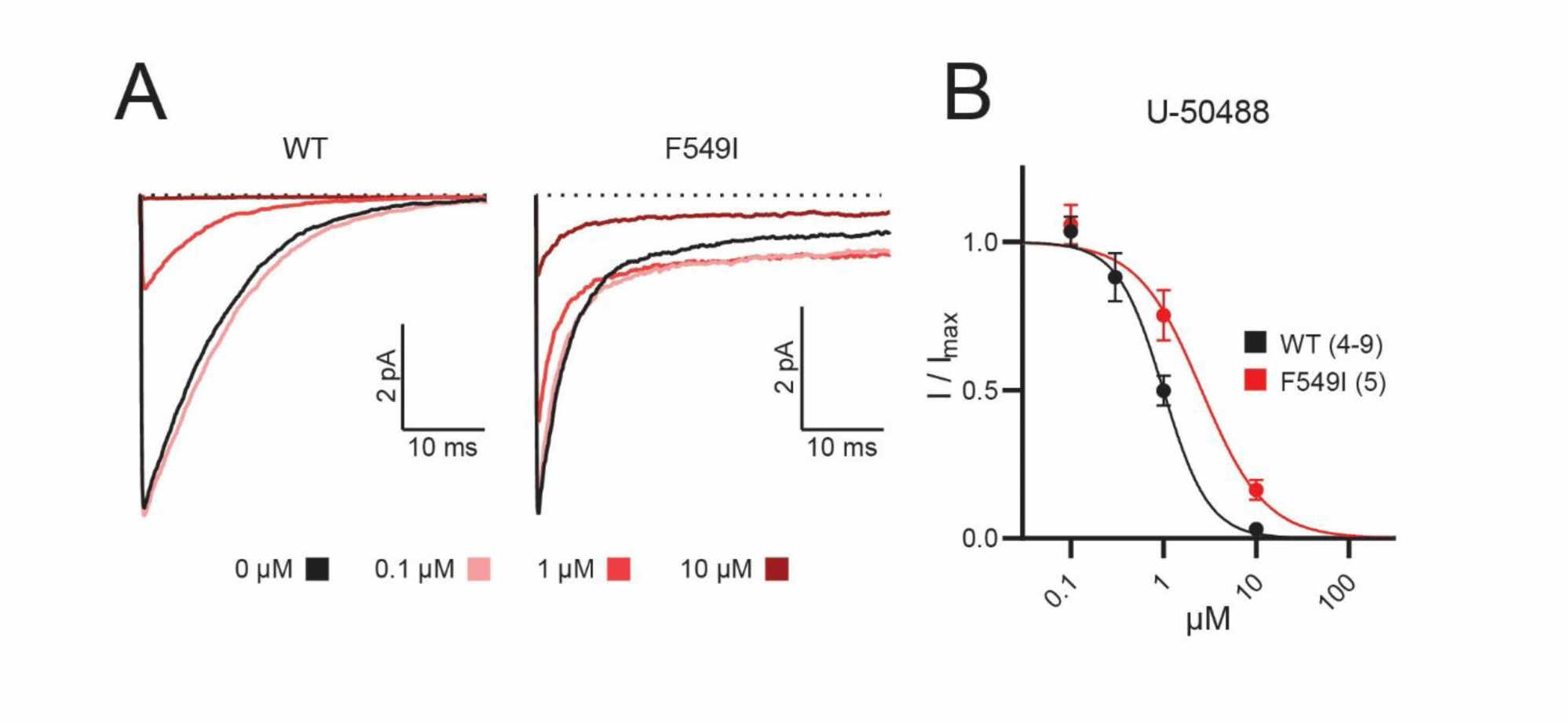
U-50488 binds to the F546 lateral fenestration receptor site. (**A**) Representative current traces from heterologously expressed PKD2L1 and F549I after treatment with U-50488. (**B**) Dose-response relationship for inhibition of PKD2L1 by U-50488 in WT and F549I channels. (n = 4-9 cells, Error = SEM)

**Figure S6.**
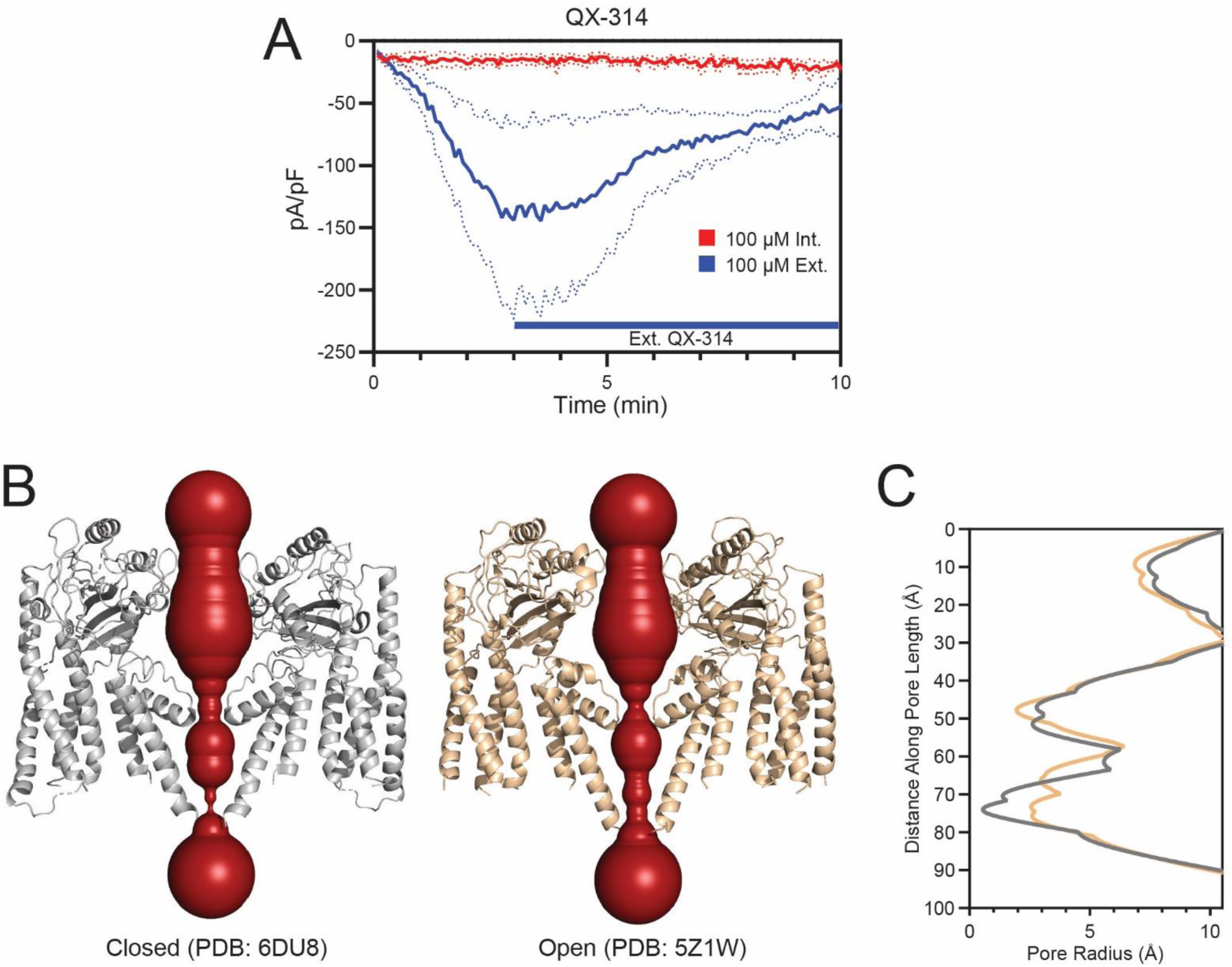
State-dependent differences in the inner gate radii restricts QX314 access to the lateral fenestration receptor. **A**) Time course of PKD2L1 tail current density during intracellular or extracellular treatment with 100 µM QX-314. (n = 3 cells, Error = SEM). (**B**) HOLE analysis of the PKD2L1 pore in its closed (left) and open (right) conformations. Red surfaces represent the pore radius through the ion-conducting pathway. (**C**) Graph of HOLE-generated pore radii down the length of the ion-conducting pathway.

**Table S1 Compounds evaluated using HTS electrophysiology.** Index of all compounds tested in HTS experiments. Index numbers (as related to Figure 1), names, known pharmacological targets, stock concentrations, solvents, and sources are included for all test articles.

**Table S2 Potency of polycystin current inhibition.** Half-maximal inhibitory concentrations (IC_50_) ± SEM, organized by test-article/ion-channel groupings. Values calculated as described in Methods. Number of cells tested for each condition are in parenthesis.

**Table S3 Docking results for screen-identified antagonists.** Output statistics of docking analyses for all screen-identified antagonists and dibucaine. Average predicted binding energy, lowest predicted binding energy, and lowest predicted K_i_ listed for all ligand/target pairs tested in AutoDock4.2.

**Table S4 Relative binding free energies of PKD2L1 mutants.** Calculated change in free energy of binding for dibucaine and U-50488 against PKD2L1 mutants, relative to WT PKD2L1. Values calculated through the relationship ΔΔG = RT ln(IC_50_Mutant/IC_50_Wildtype), where R is the gas constant and T = 298 °K.

**Table S5 Mutagenic primers used to generate F549 mutant plasmids.** Mutagenic primers used in site-directed mutagenesis experiments to generate I, L, and H mutants at the PKD2L1 F549 site. Mutant sequences highlighted in bold, underlined lettering.

## Acknowledgements

P.G.D. was supported by the National Institute of Diabetes and Digestive and Kidney Diseases (R01 DK123463-01, R01 DK131118-01) and the PKD Foundation (Research Grant).

